# Rechargeable Biomineral Induced by Sulfate Reducing Bacterium *Cupidesulfovibrio* sp. HK-II

**DOI:** 10.1101/2023.08.07.552368

**Authors:** Yui Arashi, Hiroki Mochihara, Hiroko Kubota, Kei Suzuki, Yusuke Chiba, Yutaka Kato, Toshihiro Kogure, Ryota Moriuchi, Hideo Dohra, Yosuke Tashiro, Hiroyuki Futamata

## Abstract

A black precipitate produced by a sulfate reducing bacterium *Cupidesulfovibrio* sp. strain HK was investigated with multidisciplinary methods. X-ray diffraction (XRD) analysis revealed that the black precipitate was mackinawite. Cyclic voltammetry analysis showed the obvious redox peaks, and the biogenic mackinawite exhibited rechargeable properties. XRD analyses showed that the form of the rechargeable biogenic mackinawite (RBM-II) was changed by discharge and recharge treatments: Field-emission transmission electron microscope analyses revealed that lepidocrocite and amorphous iron oxide were appeared from mackinawite on discharged condition, and the three kinds of minerals were intermingled via the rechargeable treatments. Physicochemical parameters were changed regularly under the treatments, suggesting that discharge would be occurred by iron oxidation and sulfur reduction, and *vice versa*. These results indicated that dynamics of sulfur is important key process in rechargeable mechanism, supporting that a part of mackinawite was transformed to lepidocrocite and iron oxides, and *vice versa*. Microbial fuel cells (MFCs) equipped with lactate, strain HK-II and anode including RBM-II consumed lactate even under opened circuit conditions, after which MFCs generated higher current density at re-closed circuit conditions. These results demonstrated that the biogenic mackinawite is one of rechargeable materials and would play important roles in geomicrobiological reactions and biotechnology.

## Introduction

Microbes play significantly important roles in wastewater treatment, healthy cultivation of crops, human health, elements circulation on the earth, and the like, therefore, microbes are deeply related to our existence. It is one of approaches for development of sustainable society to understand microbial functions. As a novel biotechnology using overlooked microbial functions, microbial fuel cells (MFCs) have been concerned due to generate electricity with degradation of organic matters (1–5). To date, many researchers have challenged to increase current densities and decrease internal resistance, i.e., harness structure (6−8), electrode (9, 10), and effects of anode potential on current production (11−13), whereas suggesting the limitation of modern MFCs as energy-producing devices because the problem of low current density is not dissolved completely, although the power density of MFCs has been improved.

On the other hand, studies of MFCs open simultaneously new frontiers, which are electroactive bacteria (14−17), microbial communities in anode of MFCs (18−20) and extracellular electron transfer (EET) of microbes (21−25). These studies reveal diversity of electroactive bacteria and some of EET mechanisms, which accelerates to apply for wastewater treatments (26). In addition, novel relationships between microbes and minerals have been concerned and shed light for understating overlooked survival strategy of microbes. Some bacteria produce conductive nanoparticle FeS consisting of pyrrhotite (Fe_1-x_S), mackinawite, and marcasite (FeS_2_) localized extracellularly, intracellularly and on the cell surface, which functions as an efficient extracellular electron uptake (EEU) (27) and EET (28), e.g., *Shewanella loicica* strain PV-4 produces an iron-monosulfide, and the assemblage of the cells and matured iron-monosulfide (mackinawite [Fe_1+x_S]) exhibits conductivity (28, 29). These results are expected to expand the understanding of overlooked microbial metabolisms and develop MFCs.

In our previous study, an MFC produced significantly high current density for ∼1 month (13). In current study, bacteria were isolated from the surface of anode in the MFC to analyze why the MFC exhibited high performance. An isolated bacterium, strain HK-II, induced black precipitates in the presence of sulfate and ferric iron under anaerobic conditions. Because the black precipitates looked like minerals, it was predicted that the black precipitates would have conductivity, resulting in high current density as well as other conductive minerals (28, 29). As intuition, we hypothesized that the black precipitates would have a rechargeable property, which has not been reported in the literature. If the black precipitates would be a rechargeable material, it would be a clue to develop biotechnology including MFCs and to expand the understanding of microbial ecosystems. The aims of this study were to characterize the black precipitate using material science and electrochemical techniques, and to discuss about the rechargeable mechanism and the geomicrobiological role.

## MATERIALS AND METHODS

### Isolation, incubation, and identification of bacteria

It has been reported that an MFC constructed with BE medium (9), sodium lactate as an electron donor, and lake sediment as inoculum produced a high power density of over 200 mW m^−2^ from day 168 to day 197 (13). In this study presented here, a part of the anode in the MFC was taken anaerobically in a COY chamber (COY Lab., Grass Lake, MI, USA) at day 205 when the power density was stable at 5.2 mW m^-2^. A modified BE medium (BELF medium), which was supplied with 0.1 mM Fe (III)-EDTA, was used to isolate bacteria with 0.0075% titanium (III) citrate and 20 mM sodium lactate. White, brown, gray, and black colonies were then obtained from biofilms on the surface of the anode in the MFC (13) using the roll tube method (Fig. 1A). When the gray colony was incubated in the BELF medium with 0. 5 mM Fe (III) citrate instead of 0.1 mM Fe (III)-EDTA (M-BELF medium), black precipitation was produced; however, the color of the precipitation was dark white at first before changing to black. Therefore, we tried to purify microorganisms from the culture using a six-well plate method (30) with the M-BELF medium. White and black colonies were obtained, and these colonies were purified twice using the six-well plate method. An isolated microorganism (termed strain HK-II), which formed a black colony, produced a black precipitate in the M-BELF medium (Fig. 1B). The secondary modified BE medium (termed LS medium) was used to incubate the strain HK-II to prevent production of the black precipitate, consisting of 0.5 g KH_2_PO_4_, 0.5 g KH_4_Cl, 2.5 g NaHCO_3_, 0.16 g MgCl_2_·6H_2_O, 1.0 mL SL8 solution, 1.0 mL Se/W solution, 0.15 g CaCl_2_·2H_2_O, 40 mM sodium lactate, 20 mM disodium sulfate, 0.5 mg Resazurin, 0.0075% titanium (III) citrate, and 1.0 mL vitamin solution PV1 (31). DNA was extracted and almost full-length 16S rRNA gene was amplified by PCR with the primers 5’-AGAGTTTGATCCTGGCTCAG-3’ and 5’-AAGGAGGTGATCCAG CC-3’. The nucleotide sequence of the 16S rRNA gene was analyzed using a model 377 DNA sequencer (Applied Biosystems, Foster City, CA, USA) with the accession number LC612775. The GenBank database search was conducted using the Blast version 2.11.0. A neighbor-joining tree (32) was constructed using the njplot software in ClustalW version 1.7 (Fig. S1).

**Figure 1.**
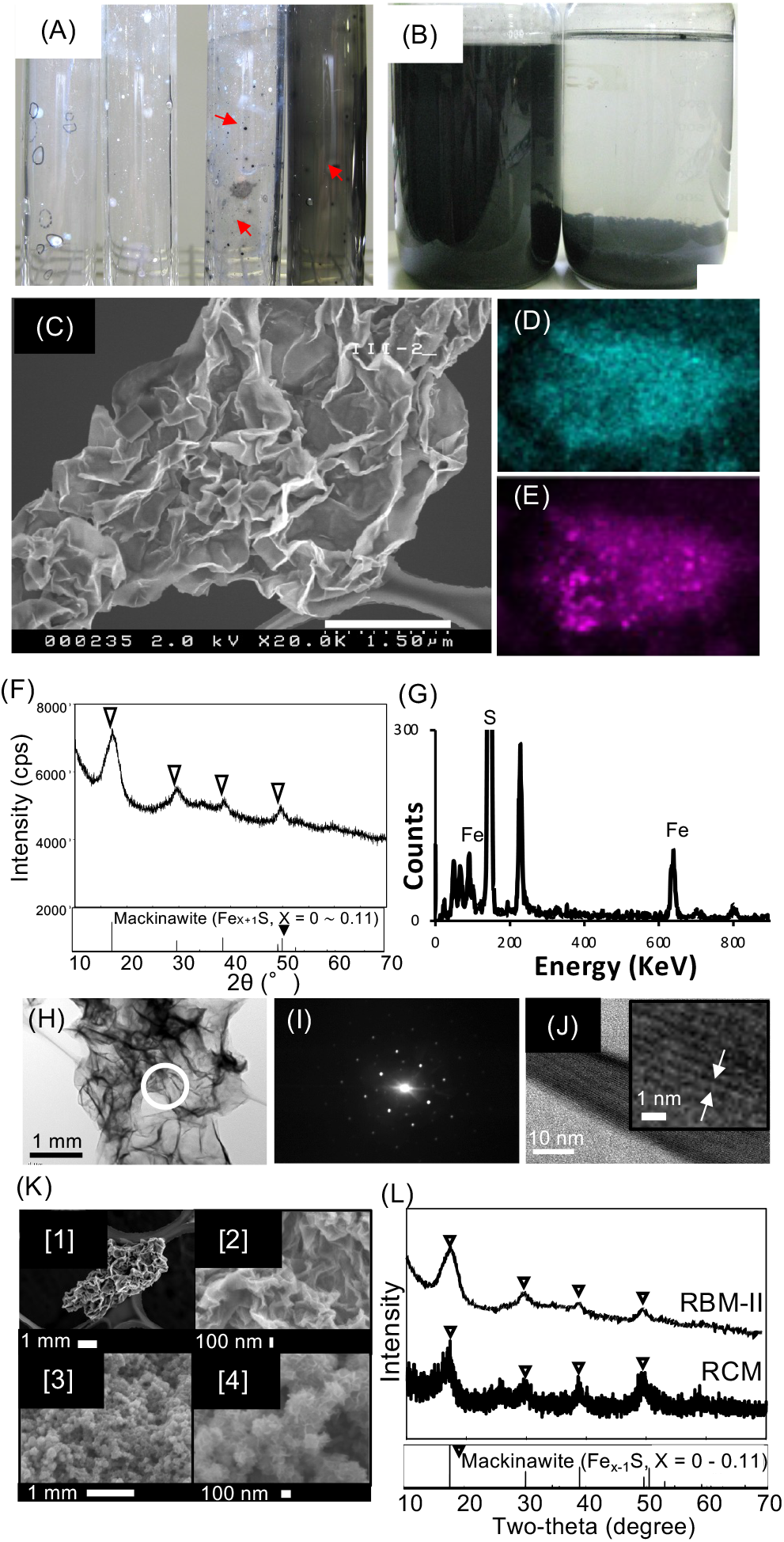
Isolation of strain HK-II and identification of black precipitates. (A) Colonies isolated from anode surface of MFC (13) with a roll tube method. Red arrows show black colored colonies. (B) Photographs show the black precipitate (RBM-II) produced by strain HK-II. The left bottle is just after agitation. The right bottle is after static condition for 30 min. (C) Scanning electron microscopy (SEM) image of RBM-II. The scalebar indicates 1.50 μm, (D) Iron mapping by energy dispersive X-ray (EDX) analysis of RBM-II, (E) sulfur mapping by EDX analysis, (F) X-ray diffraction (XRD) analysis profile of RBM-II, (G) EDX analysis profile of RBM-II, and (H) transmission electron microscopy (TEM) image of RBM-II. The scalebar indicates 1 μm. The white circle shows the area for diffraction analysis in (I), (J) High-resolution TEM images showing mackinawite lattice fringes on RBM-II. The scalebar indicates 10 nm. The small picture in (J) shows the distance between layers was 5 Å, which corresponds with the reference values for pure mackinawite (83). (K) SEM observation of RBM-II and RCM. (K_1): RBM-II. The white scalebar indicates 1 μm, (K_2): RBM-II. The scalebar indicates 100 nm, (K_3): RCM. The scalebar indicates 1 μm, (K_.4): RCM. The scalebar indicates 100 nm. (L) XRD analyses of RBM-II and RCM. The triangle “ ” indicates representative peaks of mackinawite shown in the Powder diffraction file #86-0389. RBM-II: rechargeable biomineral induced by strain HK-II; RCM: rechargeable chemically synthesized mackinawite.

### Electrochemical analysis of the black precipitate produced by strain HK-II

The strain HK-II was incubated in the M-BELF medium and the produced black precipitate was collected by filtration with a membrane filter (pore size of 0.1 μm, Advantec, Tokyo, Japan) and was washed with M-BELF medium, excluding sodium lactate, MgSO_4_·7H_2_O, and Fe (III) citrate, under anaerobic conditions in a COY chamber (COY Lab.). A part of the black precipitate collected on the filter was suspended in 2.0 mL M-BELF medium and then the suspended black precipitate was attached to a carbon felt using a suction pump. The carbon felt attached to the black precipitate was set in a single-chamber MFC (volume 7 cm^3^) with a three-electrode system (HX-R6 working, counter, and Ag/AgCl reference electrodes; Hokuto Denko, Tokyo, Japan), and the MFC was filled with M-BELF medium, excluding sodium lactate, MgSO_4_·7H_2_O, and Fe (III) citrate. Low-scan cyclic voltammetry (LSCV) was performed at a scan rate of 1 mV s^−1^ between −1.1 V and 0.8 V vs SHE. A voltage was chosen for charging the black precipitate according to the result of LSCV, in which the potential (−0.55 V) was more negative than the lowest potential at the reductive peak. The voltage was monitored by connecting potentiostat (HA-151B; Hokuto Denko) with a data logger GL200A (Graphtec, Yokohama, Japan). When the voltage plateaued, the charge was defined as complete. A discharge from the black precipitate was performed, in which the voltage was monitored between the working electrode connected to the anode (the carbon felt attached to the black precipitate) and the counter electrode was connected to the cathode using 10 Ω as the external resistance. When the voltage was almost stable (∼ 0 V), the discharge was defined as complete. The capacity was calculated using the following formulas: 𝑉 = 𝐼𝑅 and 𝐶 = 𝐼𝑇, where *V* is the voltage (in V), *I* is the current (in A), *R* is the resistance (in Ω), *C* is Coulomb (in C), and *T* is time (in s). Physicochemical parameters in the single-chamber MFC were monitored on the rechargeable treatments, and the pH was measured using a liquid sample with a pH meter (twinpH or B-212; Horiba, Kyoto, Japan). Sulfate was measured by high-performance liquid chromatography (HPLC) equipped with an Shodex IC NI-424 column (100 × 4.6 mm) (Showa Denko, Tokyo, Japan) and a conductivity detector (Shodex CD-5; Showa Denko). Sulfide was measured using the methylene blue colorimetric method. The proportion of sulfur to iron and the percentage of oxygen and phosphorus were calculated using the results of the energy dispersive X-ray spectrometry analyses.

### Scanning electron microscopy

SEM observations and qualitative element analyses of the black precipitate were performed using a scanning electron microscope (S-4500; Hitachi, Tokyo, Japan) equipped with an energy dispersive X-ray spectrometer. The sample (∼20 μL) of the culture was dispersed onto a carbon-coated copper grid and the grid was mounted on a standard aluminum stub. The applied accelerating voltage was 15 kV for EDX analyses.

### Energy dispersive X-ray spectrometry

The strain HK-II was incubated in BELF medium for 2 weeks. The black precipitate was collected by filtration with a membrane filter (pore size of 0.1 μm, Advantec) under anaerobic conditions in a COY chamber. The black precipitate on the membrane was subjected to EDX analysis (Miniscope TM3000; Hitachi).

### X-ray diffraction analysis

The strain HK-II was incubated in the BELF medium for 9 days and the produced RBM was then collected by filtration with a membrane filter (pore size of 0.1 μm, Advantec) under anaerobic conditions in a COY chamber and stored under anaerobic conditions until analysis. XRD analyses were conducted with Cu-*K*α radiation (wavelength 1.5618 Å) at a scan rate of 5° per min from 10° to 70° at 0.02° steps in 2*θ* at 1.2 kW of output using X-ray analysis instrumentation (RINT2200; Rigaku., Tokyo, Japan). The average diameters of the crystalline domains were determined from the XRD patterns using the Scherrer equation: 𝐿 = 𝐾𝜆(𝛽𝑐𝑜𝑠𝜃)^!"^ (33), where *L* is the average diameter of the domain, *K* is the Scherrer constant (0.91), *λ* is the wavelength of the applied X-rays (1.5418 Å), *β* is the full width (in radians) at half maximum of the peak, and *θ* is the angle at the position of the peak.

### MFC configuration and operation

Mediator-less and air-cathode MFCs were constructed to evaluate whether RBM-II (the black precipitate was termed RBM-II) is capable of charging electrons produced by microorganisms. A carbon paper electroplated with platinum (0.5 mg cm^−2^) on one side was used as the cathode electrode (Chemix, Sagamihara, Japan), providing a total projected cathode surface area (on one side) of 3.06 cm^2^ (a window of 1.75 cm × 1.75 cm set on the outside of the MFC). A proton exchange membrane (Nafion 117, Dupont, Wilmington, DE, USA) was placed between the anode and the cathode. Graphite felts (SOHGOH-C., Yokohama, Japan) were used as the anode (4 cm × 4 cm × 0.5 cm) and were packed in the anode chamber (50 mL capacity) to provide a projected anode surface area of 40 cm^2^. RBM-II was produced by strain HK-II in BE medium (33) and was collected on a membrane filter (0.1 μm, OmniporeFM membrane filter; Merck Millipore, Burlington, MA, USA) using a suction pump, and RBM-II was washed three times with autocleaved anaerobic dH_2_O. Before setting the anode in the anode chamber, a slit (∼2.5 cm × 3 cm) was set in the anode by cutting with an autocleaved knife, and then RBM-II was added into the slit (anode-RBM-II electrode). The slit was sewed using a fishing line. Three MFCs equipped with the anode-RBM-II electrode were constructed: RBM-MFC-1, RBM-MFC-2, and RBM-MFC-3, in which RBM-II added were 0.24 g, 0.25 g, and 0.33 g, respectively. RBM-II was not added to all of control-MFC-1, -2, and -3. Strain HK-II was incubated in NB medium, and the cells were then collected by centrifugation (4000 ×*g* for 40 min) after sulfate was almost completely consumed by the sulfate-reduction process of strain HK-II (detection limit was 2.5 μM). The cells were washed three times with a modified NB medium (AE medium) containing 0.16 g MgCl_2_·6H_2_O instead of 2.0 g MgSO_4_·7H_2_O per liter. The cells were resuspended in AE medium and inoculated at OD_600 nm_ of 0.2 into the anode chamber of all MFCs. Sodium lactate 30 mM was added as an electron donor. The external resistance (51 Ω) was connected between the anode and the cathode using a platinum wire. All the procedures were performed in a COY chamber under anaerobic conditions. All MFC voltages were recorded every 5 min across 51 Ω resistance using a data logger. All MFCs were run under semi-batch conditions at 25 ℃, and fresh sodium lactate was added at 30 mM when lactate was consumed by sulfate-reduction process of strain HK-II. Because the voltage of all MFCs was stable after sodium lactate was added to all MFCs on day 39, the circuit was opened to charge electrons into RBM-II for 4 h, and then closed again. If RBM-II is charged, the current density after the re-closed circuit is higher than that before the opened circuit. The culture solution in the anode chamber was sampled adequately, and the OD_600nm_ and concentration of organic acids was measured with spectrophotometer (UV-1800; Shimadzu, Kyoto, Japan) and HPLC (GL-7410 and 7450, GL Science, Tokyo, Japan), respectively. The charged capacity was calculated using the following formula, 𝐶 = 𝐼𝑇, where *C* is the Coulomb (in C), *I* is the current (in A), and *T* is time (in s). *T* was defined as the time (s) when the circuit was closed to the time when the current density decreased to the level before the circuit was opened. Coulombic efficiency was obtained by calculating the ratio of the charged capacity per theoretical amount of coulombs produced by consuming organic acids during the open circuit (4 h).

### Chemical synthesis of mackinawite

Mackinawite was synthesized chemically by mixing the same volume solutions of 100 mM Na2S·9H2O and 100 mM Fe(SO4)2 (NH4)2·6H2O under anaerobic conditions (34). Before mixing these solutions, the headspace gas in a bottle containing Na2S·9H2O solution was exchanged twice with nitrogen gas for 20 min, and the Fe(SO4)2 (NH4) 2·6H2O solution was degassed for 60 min and purged with high-purity nitrogen gas for 40 min twice. In additional to RBM-II, CSM was analyzed by XRD, XPS, SEM, and electrochemical techniques according to the methods described above.

### Chemical analyses

Liquid samples were collected from all the MFCs. These liquid samples were also filtered (Millipore LG [pore size: 0.2 μm, diameter: 13 mm]; Merk Millipore) for quantification of organic acids using HPLC equipped with Shodex RSpak KC-811 column (300 × 8.0 mm) (Showa Denko) and a UV detector. The column heater was set to 50°C, samples were eluted using 0.1% H3PO4 solution delivered at 1.0 mL min^−1^, and elutes were monitored at 210 nm. Formate, pyruvate, lactate, butyrate, and acetate were identified according to their retention times, and concentrations were determined by comparing the peak area with that of the cognate standard sample. The pH, sulfate, and sulfide concentrations were measured as described above.

## RESULTS

### Material characterization and scanning electron microscopy (SEM) observation of the black precipitate and chemically synthesized mackinawites

The black precipitate was induced by an isolated bacterium (Fig. 1A and B). SEM observations showed that the black precipitate was thin and frilled (Fig. 1C, Fig.1K [1] and [2]). Energy dispersive X-ray (EDX) analysis showed that the black precipitate consisted mainly of iron and sulfur (Fig. 1D, E, G). X-ray diffraction (XRD) analysis revealed that the black precipitate was crystalline, and peaks at 2 *θ* = 17.61°, 30.09°, 38.99°, and 49.55° corresponding to the [001], [101], [111], and [200] mackinawite, respectively (powder diffraction file #86-0389) (Fig. 1F). XRD analysis revealed that the average crystal size of the black precipitate was estimated to be 9.2 ± 1.5 nm (Table S1). The diffraction pattern confirmed that the black precipitate was mackinawite (Fig. 1H and I) and crystal layered structure was formed with a distance of 5.03 Å between layers (Fig. 1J). These results demonstrated that the black precipitate is biogenic mackinawite (Fe1+*X*S, *x* = 0∼0.11).

As a control, chemically synthesized mackinawite (CSM) was prepared according to a previous method (34). SEM observations showed that the morphology of CSM was an aggregate of particles with diameters of 100∼200 nm with a frilled form on the surface of the particles (Fig. 1K.3 and 4). As expected, XRD analysis revealed that CSM was crystalline, and peaks at 2 *θ* = 17.61°, 30.09°, 38.99°, and 49.55° corresponding to (001), (101), (111), and (200) (Fig. 1L) of mackinawite, respectively (powder diffraction file #86-0389), corroborating that CSM was chemically synthesized mackinawite (Fe1+xS, x = 0∼0.11) as previously reported (34). XRD analysis revealed that the average crystal size of CSM was estimated to be 10 ± 0.9 nm (Table S1).

### Electrochemical and rechargeable properties of black precipitate and CSM

Cyclic voltammetry (CV) analyses were conducted to electrochemically characterize the black precipitate and CSM (Fig. 2A and B). The black precipitate showed oxidative peak potentials at 0.24 V (vs standard hydrogen electrode [SHE] on 1^st^ cycle) and 0.17 V (vs SHE on 2^nd^ cycle), whereas the reductive peak potentials were −0.50 V (vs SHE on 1^st^ cycle) and −0.48 V (vs SHE on 2^nd^ cycle) (Fig. 2A). The CSM showed that the oxidative and reductive peak potentials were 0.34 V (vs SHE on 1^st^ and 2^nd^ cycles) and −0.64 V (vs SHE on 1^st^ and 2^nd^ cycles), respectively (Fig. 2B). Since these data indicated that the black precipitate and CSM had redox sites, the rechargeable properties of the black precipitate and CSM were investigated. The charge and discharge capacitance of the black precipitate was 160 ± 35 mAh g^−1^ and 53 ± 14 mAh g^−1^, respectively, whereas the charge and discharge capacitance of the CSM was 250 ± 57 mAh g^−1^ and 94 ± 30 mAh g^−1^, respectively (Fig. 2C). The black precipitate was named RBM-II (rechargeable bio-mineral induced by strain HK-II).

**Figure 2.**
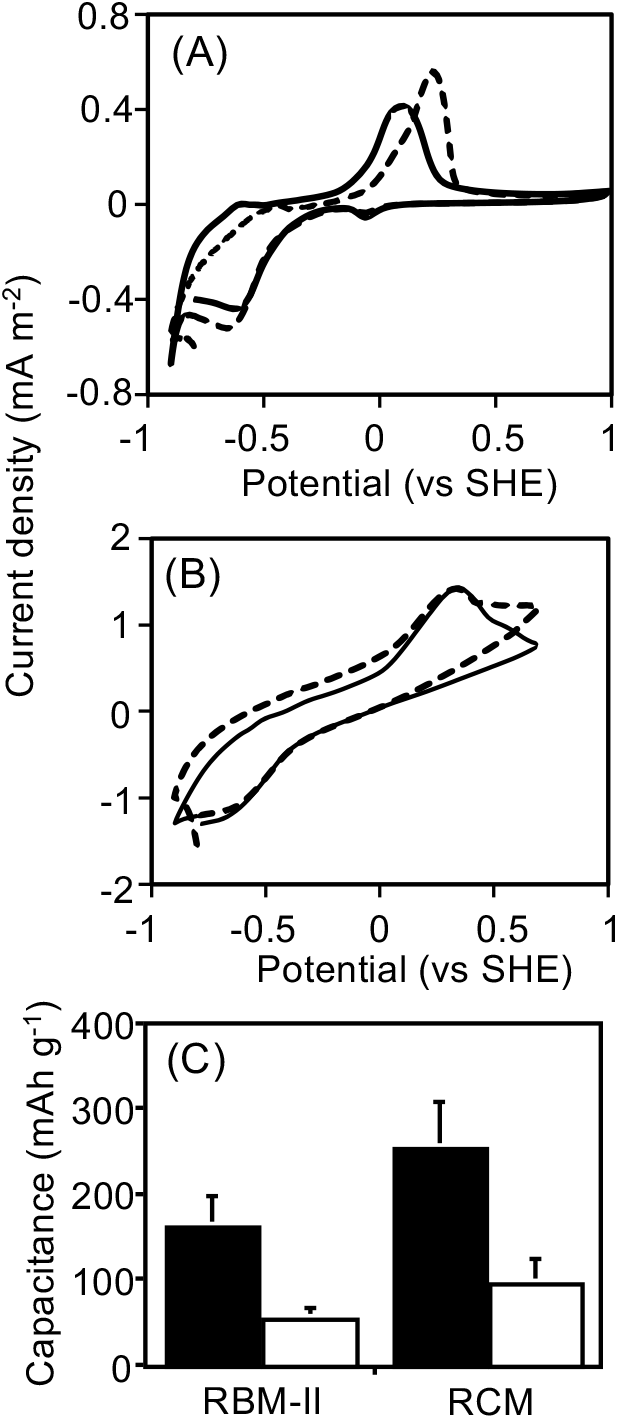
XRD analyses and electrochemical analyses of RBM-II and RCM. (A) Cyclic voltammetry profiles of RBM-II. The broken and solid lines show first and second cycles, respectively. (B) Cyclic voltammetry profiles of RCM. The broken and solid lines show first and second cycles, respectively. (C) Charge and discharge capacitys of RBM-II and RCM are shown in black and white bars, respectively. SEM: scanning electron microscopy; XRD: X-ray diffraction; RBM-II: rechargeable biomineral produced by strain HK-II; RCM: rechargeable chemically synthesized mackinawite.

### pH and concentrations of sulfide and elements in RBM-II and CSM under charged and discharged conditions

The pH in RBM-II suspension decreased from 6.9 ± 0.22 in the 2^nd^ discharged condition to 5.3 ± 1.1 on the 2^nd^ charged condition (Fig. 3A-1). After that, the pH of the suspension did not decrease under discharged conditions but decreased in charged conditions. The sulfide concentration in the RBM-II suspension increased to 0.18 ± 0.044 μM in the 2^nd^ discharged condition and decreased to 0.12 ± 0.025 μM in the 2^nd^ charged condition. The sulfide concentration increased and decreased slightly under the 4^th^ and 6^th^ discharged and charged conditions, respectively (Fig. 3A-2). Furthermore, sulfate was not detected in the RBM-II suspensions. The proportion of sulfur to iron changed periodically corresponding to the charged and discharged conditions. The proportions were 0.83 ± 0.046 on discharge and 1.0 ± 0.053 on charge (Fig. 3A-3). The percentages of oxygen and phosphorus changed periodically corresponding to the charged and discharged conditions. The percentage of oxygen in RBM-II was 23% ± 3.7% in discharge and 15% ± 2.1% in charge (Fig. 3A-4), whereas the percentage of phosphorus in RBM-II was 0.42% ± 0.042% in discharge and 0.083% ± 0.037% in charge (Fig. 3A-5).

**Figure 3.**
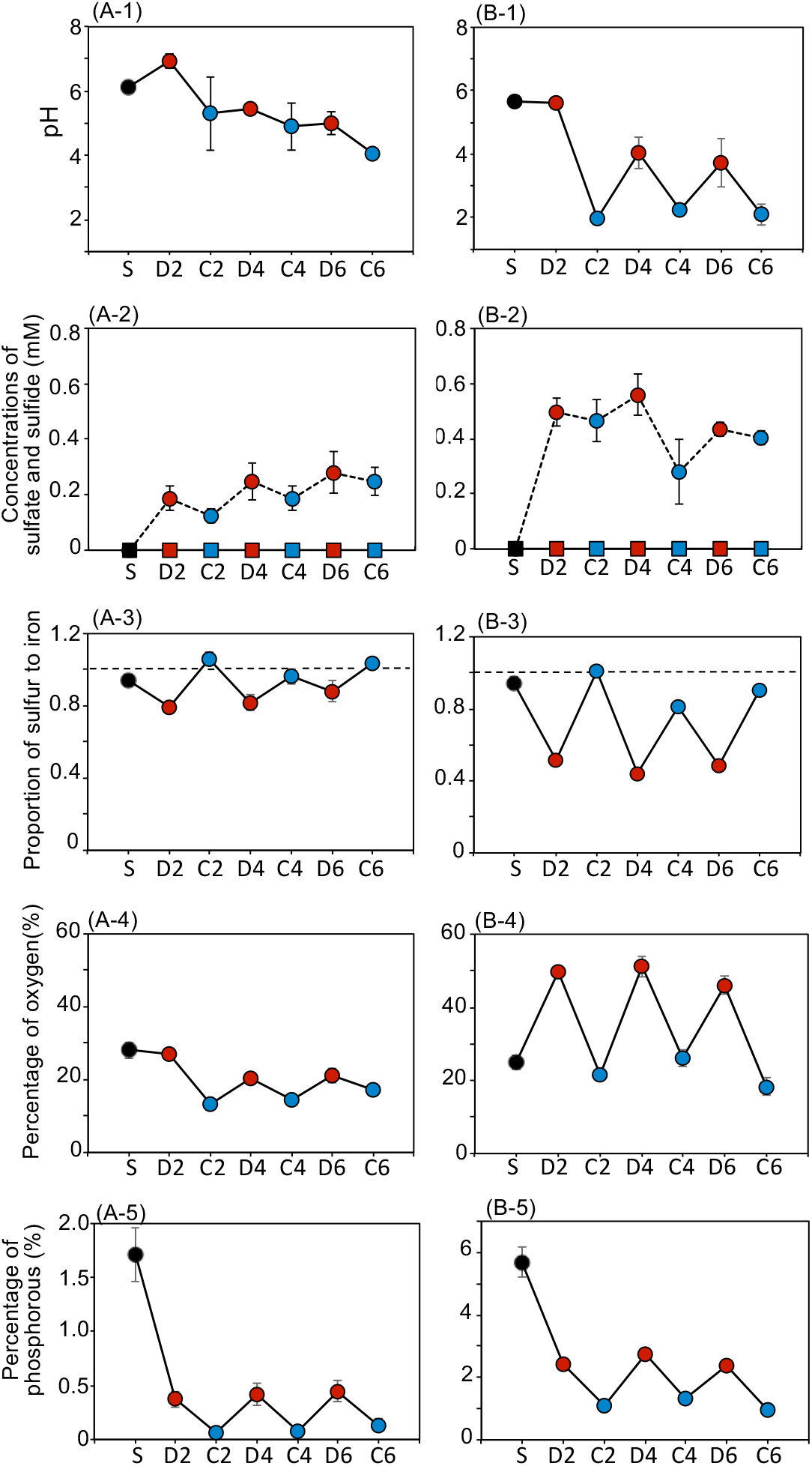
Physicochemical parameters of RBM-II and RCM under rechargeable treatments. (A-1_-5), RBM-II and (B-1_-5), RCM. (A-1)/(B-1) pH, (A-2)/(B-2) Concentrations of sulfide (broken line) and sulfate (solid line). (A-3)/(B-3) Proportion of sulfur to iron. The broken line denotes that the proportion of sulfur to iron is 1.0. (A-4)/(B-4) Percentage of oxygen. (A-5)/(B-5) The percentage of phosphorous. S: initial sample after RMB-II was produced by strain HK-II or RCM was synthesized chemically; D: discharged sample; C: charged sample, the number beside “D” and “C” denotes the number of discharge and charge cycles. RBM-II: rechargeable biomineral produced by strain HK-II; RCM: rechargeable chemically synthesized mackinawite.

**Figure 4.**
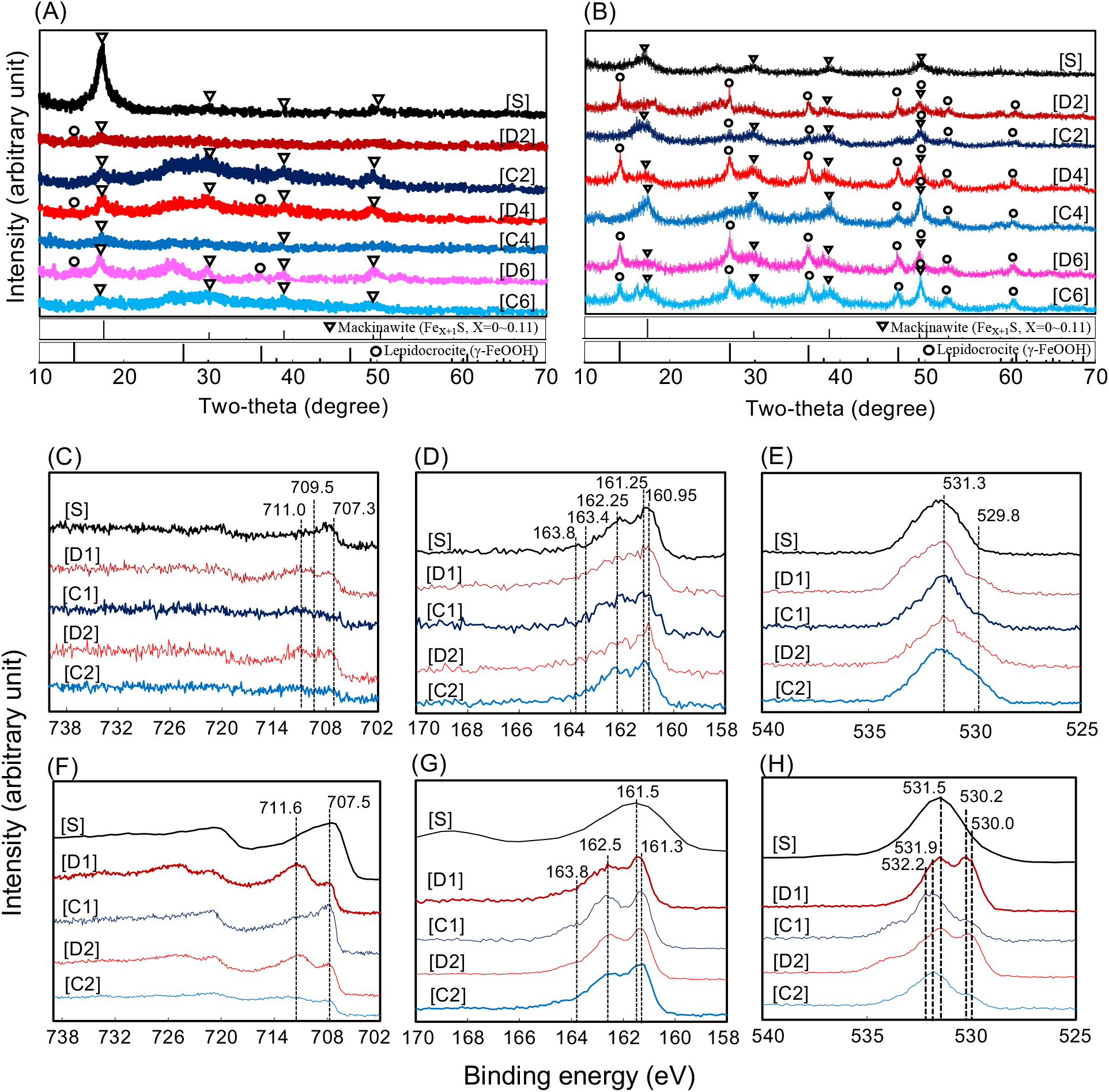
XRD and narrow scans of XPS analyses of RBM-II and RCM under discharge and charge conditions. (A) XRD patterns of RBM-II, (B) XRD patterns of RCM. The triangle “ ” indicates representative peaks of mackinawite shown in the Powder diffraction file #86-0389. The circle “ ” indicates representative peaks of lepidocrocite shown in the Powder diffraction file #74-1877. (C) Fe(2p^3/2^), (D) S(2p), and (E) O(1s) of RBM-II, respectively. (F) Fe(2p^3/2^), (G) S(2p), and (H) O(1s) of RCM, respectively. [S]: initial sample after RMB-II was produced by strain HK-II or when RCM was synthesized chemically; [D]: discharged sample; [C]: charged sample, the number beside “D” and “C” denotes the number of discharge and charge cycles. Binding energies for Fe(2p^3/2^), S(2p), and O(1s) peaks were described in the figure with dush lines and were shown in SI Appendix, Table S2. XRD: X-ray diffraction, XPS: X-ray photoelectron spectroscopy; RBM-II: rechargeable biomineral produced by strain HK-II; RCM: rechargeable chemically synthesized mackinawite.

**Figure 5.**
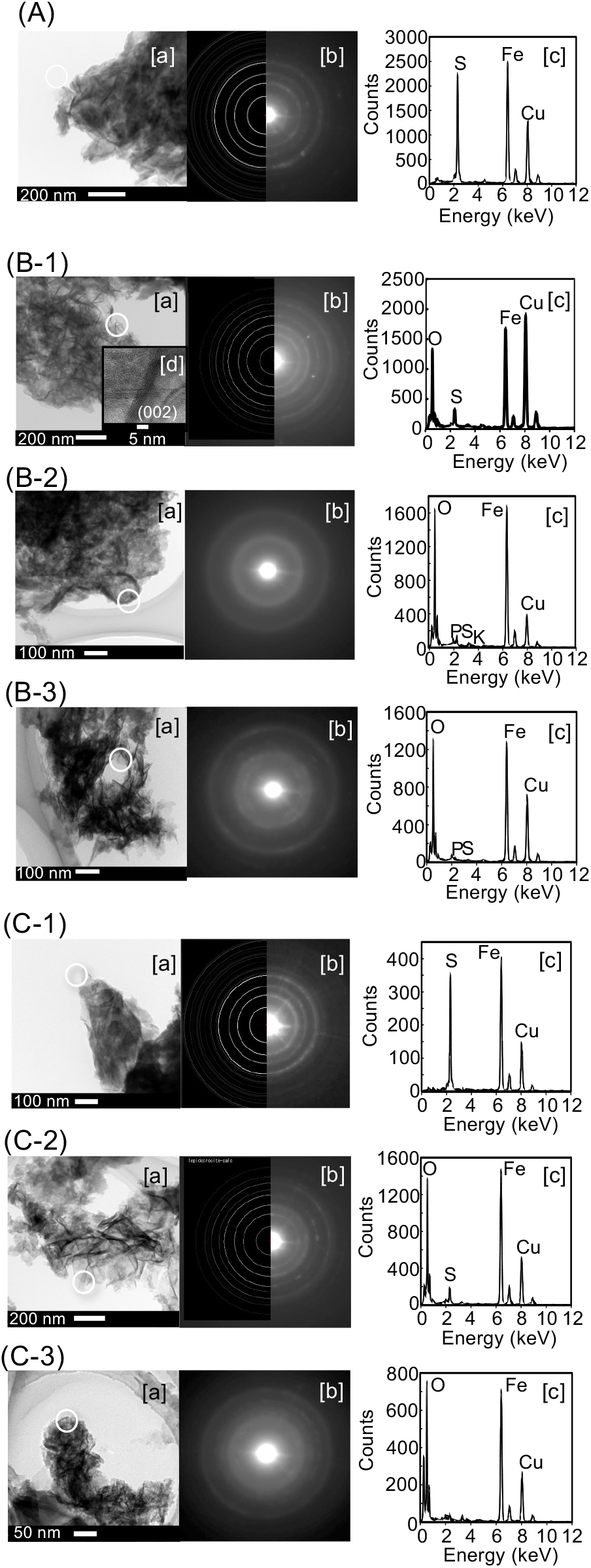
Form changes of RBM-II confirmed by FE-TEM. (A) RBM induced by strain HK-II, (B) The 2^nd^ discharged RBM-II. Different three positions were analyzed (B-1, -2, and -3), (C) The 2^nd^ charged RBM-II. Different three positions were analyzed (C-1, -2, and -3), [a] High-resolution TEM images, [b] Electron diffraction patterns at a selected area, [c] EDX spectra, [d] in B-1-a: Displaying characteristic (002) layers of lepidocrocite. A white bar is size marker. A white circular shows the area for diffraction analysis.

On the other hand, the pH in the CSM suspension decreased from 5.6, in the 2^nd^ discharged condition, to 1.8 ± 0.20 in the 2^nd^ charged condition. After that, the pH in the suspension changed regularly from 4.0 ± 1.5 on discharged conditions to 2.6 ± 1.1 on charge conditions (Fig. 3B-1). The sulfide concentration in the CSM-suspension increased by 0.50 ± 0.051 μM under the 2^nd^ discharged condition and decreased by 0.47 ± 0.076 μM under the 2^nd^ charged condition (Fig. 3B-2), after which it increased and decreased slightly under discharged and charged conditions, respectively. Furthermore, sulfate was not detected in the CSM suspension during this experiment. The proportion of sulfur to iron changed periodically corresponding to the charged and discharged conditions, which were 0.48 ± 0.036 on discharge and 0.91 ± 0.10 on charge (Fig. 3B-3). Moreover, percentages of oxygen and phosphorus changed periodically corresponding to the charged and discharged conditions. The percentage of oxygen in CSM was 49% ± 2.6% in discharged and 22% ± 4.0% in charged conditions (Fig. 3B-4), whereas the percentage of phosphorus in CSM was 2.5% ± 0.17% in discharge and 1.1% ± 0.18% in charge (Fig.3B-5).

### Form-changes in RBM-II and CSM

XRD analyses were conducted to investigate the form-changes in RBM-II and CSM corresponding to the discharged and charged conditions (Fig. 4A and B) because the surface structure of RBM-II and CSM was changed by discharge and charge treatments (Fig. S2 and S3). The 2^nd^ discharged CSMs had representative peaks of lepidocrocite (Power diffraction file#74-1877) (Fig. 4B-[D2]), whereas the 4^th^ charged CSMs had those of mackinawite (Power diffraction file#86-0389) with disappear of some representative peaks of lepidocrocite (Fig. 4B-[C4]). Both representative peaks of lepidocrocite and mackinawite were observed under 6^th^ discharged and charged conditions (Fig. 4B-[D6, C6]). By comparing with CSM, the XRD results of RMB-II were ambiguous because the representative peaks of lepidocrocite were unclear under discharged conditions (Fig. 4A).

X-ray photoelectron spectroscopy (XPS) analyses were conducted to confirm whether RBM-II and CSM changed in accordance with the charge and discharge conditions in detail. In the case of CSM, the binding energy of Fe(2P^3/2^) at 711.6 eV corresponding to the Fe(III)-O band (35) was higher than that of 707.3 eV and 707.5 eV corresponding to the Fe(II)-S bond (36, 37) on discharged conditions (Fig. 4F-[D1] and [D2]), and vice versa (Fig. 4F-[S], [C1] and [C2]). In RBM-II, the binding energy of Fe(2P^3/2^) at 711.0 eV corresponding to Fe(III)-O (38) was higher than that of 707.3 eV corresponding to the Fe(II)-S bond (37) on discharge conditions (Fig. 4C-[D1] and [D2]), and vice versa (Fig. 4C-[S], and [C2]). The binding energy of Fe(2P^3/2^) at 709.5 eV corresponding to Fe(II)-O (36) increased and decreased after discharge and charge treatments, respectively (Fig. 4C).

One broad peak of the binding energy of S(2p) at around 161.5 eV was observed in the initial CSM (Fig. 4G-[S]), whereas the two main peaks at 161.3 eV and 162.5 eV corresponding to monosulfide (39) and disulfide (40), respectively, were always observed on discharge and charge conditions (Fig. 4G). In RBM-II as well as CSM, the two main peaks of binding energy of S(2p) at 160.95 eV and 162.25 eV corresponding to monosulfide (36) and disulfide (39), respectively, were observed under all conditions with the exception of the 1^st^ discharged condition (Fig. 4D).

The binding energy of O(1s) at 531.5 eV corresponding to the OH^−^ component (36) was observed in the initial CSM, and was one of the major contributors to discharged conditions (Fig. 4H-[S], [D1], and [D2]). A new broad peak in binding energy of O(1s) at 530.0 eV (41) and 530.2 eV (35) corresponding to O^2−^ were observed under discharged conditions (Fig. 4H-[D1] and [D2]). The broad peak disappeared after charge and only one broad peak of binding energy at around 531.9 eV and 532.2 eV was observed but unidentified (Fig. 4H-[C1] and [C2]). In RBM-II, the binding energy of O(1s) at 531.3 eV corresponding to OH^−^ component (42) was a major contribution in RBM-II under all conditions (Fig. 4E). The binding energies for Fe(2P3/2), S(2p), and O(1s) peaks are listed in Table S2.

FE-TEM analyses were conducted to investigate the form-change of RBM-II in detail. TEM observation showed that RBM-II was like film form (Fig. 5A-[a]). Electron diffraction pattern and EDX analyses showed that the charged RBM-II was mackinawite (Fig. 5A-[b] and –[c]). After the 2^nd^ discharge, a scaly form was observed (Fig. 5B-[a]) and the occurrence of lepidocrocite was confirmed in selected area by electron diffraction pattern and EDX analyses (Fig. 5(B-1)-[b] and –[c]). High-resolution TEM revealed that the nanocrystal lepidocrocite consisted with the (020) plane with spacing of 0.64 Å was observed (Fig. 5(B-1)-[d]), whereas amorphas oxidized irons were also observed (Fig. 5(B-2) and 5(B-3)). After the 2^nd^ charge, electron diffraction pattern and EDX analyses demonstrated that RBM-II was mackinawite (Fig.5(C-1)), whereas lepidocrocite and an amorphous iron oxide were observed in another selected area (Fig. 5(C-2) and 5(C-3), respectively). These results indicated that RBM-II was consisted of mackinawite, lepidocrocite, and amorphous iron oxide after the 2^nd^ discharge treatment.

### Charge and discharge of RBM-II in the MFCs

In control-MFCs, the consumption of initially added lactate started around day 10 and was almost consumed at day 27 in control-MFC1 and at day 37 in control-MFC2 and control-MFC3 (Fig. S4 D-F). The average current density of control-MFCs was 5.8 ± 2.8 mA m^−2^ during the initial lactate consumption. On the other hand, in RBM-MFCs, high current densities were observed in all RBM-MFCs in the initial phase (from day 0 to day 1) without lactate consumption, whereas none of the control-MFCs produced current (Fig. 6A). A positive correlation was observed between the initial current density and the amount of RBM-II added to the anode, suggesting that the initial current was produced from RBM-II. the consumption of initially added lactate started around day 10 and was almost consumed at day 18 (Fig. S4 A-C). The average current density of RBM-MFCs was 18 ± 9.9 mA m^−2^ during the first lactate consumption. The lactate consuming rate and the current density increased and were stable in all MFCs after the consumption of initial lactate (Fig. 6A).

**Figure 6.**
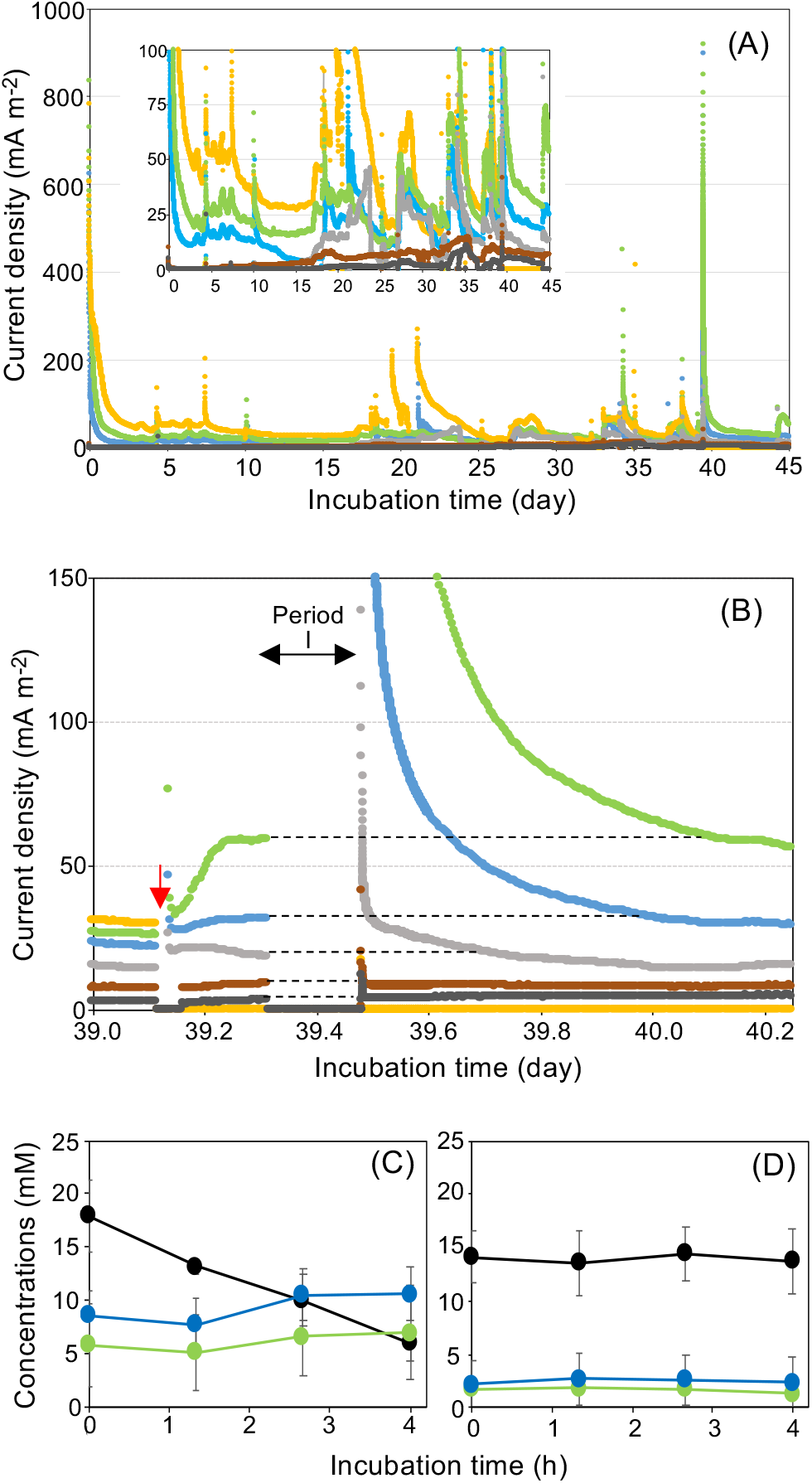
Current productions of MFCs equipped with RBM. (A) and (B) The current density of RBM-MFCs and control-MFCs. Blue, green, and orange color lines show the current density of RBM-MFC1, 2, and 3, respectively. Gray, brown, and black color lines show the current density of control-MFC1, 2, and 3, respectively. The small figure in figure A shows the difference of current density between RBM-MFCs and control-MFCs. (B) The current density of all MFCs under closed and opened circuit conditions. The red arrow shows the time of lactate addition in RBM-MFCs. The red arrow denotes that when culture solutions were collected from all MFCs, the datalogger was removed from MFCs so that the current densities were zero. The period I shows the discharging time when the circuit was opened in all MFCs. The electrode wire of RBM-MFC3 was broken when connecting wires of anode and cathode after period I. Therefore, the current from RBM-MFC3 was not detected (orange line). (C) The concentrations of organic acids during open circuit (Period I) in RBM-MFCs. Black circule: lactate; Blue circule: acetate; Green circle: propionate. (D) The concentrations of organic acids during open circuit (Period I) in control-MFCs.Black circule: lactate; Blue circle: acetate; Green circle: propionate.

The current density became stable at 31 ± 2.7 mA m^−2^ in RBM-MFC1 and 58 ± 2.1 mA m^−2^ in RBM-MFC2 after the addition of fresh lactate at day 39, and then the circuit was opened for 4 h (Period I shown in Fig. 6B) to investigate whether RBM-II was recharged by bacterial activity (Fig. 6B). Unfortunately, the RBM-MFC3 was broken so that its current could not be measured during the experiment. After re-closing the circuit, the current production from RMB-MFC1 and RBM-MFC2 was observed: the average current produced from the charged RBM-II was 120 ± 90 mA m^−2^ and 250 ± 140 mA m^−2^ in RBM-MFC1 and RBM-MFC2, respectively (Fig. 6B). It took 13.6 h and 22.9 h in RBM-MFC1 and RBM-MFC2, respectively, when the current density became to the similar level at before opening the circuit (Fig. 6B).

Concentrations of organic acids were monitored during Period I (Fig. 6C and D). Concentrations of lactate, acetate, and propionate were stable in all control-MFCs (Fig. 6D), while lactate concentration decreased, and acetate and propionate concentrations increased in all RBM-MFCs (Fig. 6C). Theoretical coulombs produced from organic acids were estimated to be approximately 1,060 kC and 450 kC in RBM-MFC1 and RBM-MFC2, respectively. The charged capacity of RBM-MFC1 and RBM-MFC2 were 170 and 390 kC, respectively. Coulombic efficiency for the theoretical coulomb of the charged capacity produced by organic acids was 16% and 86% in RBM-MFC1 and RBM-MFC2, respectively.

## DISCUSSION

This study demonstrated that a mineral induced by *Cupidesulfovibrio* sp. strain HK-II under sulfate reducing condition with ferric iron has rechargeable properties. The rechargeable property was maintained with regular change in physiochemical parameters (Fig. 2−5), which were clues to unveiling the rechargeable reactions of RBM-II. From these results, when RBM-II would be completely form change on discharge and charge conditions, simply reactions were predicted as follows (Fig. S5-A): about a discharge reaction from mackinawite to lepidocrocite, 4𝐹𝑒𝑆 + 8𝐻_2_𝑂 → 4γ-𝐹𝑒𝑂𝑂𝐻 + 4𝐻_2_𝑆 + 4𝐻^+^ + 4𝑒^-^ (reaction on anode [1]), 4𝐻^+^ + 𝑂_2_ + 4𝑒^-^ → 2𝐻_2_𝑂 (reaction on cathode [2]), and total reaction was as follows: 4𝐹𝑒𝑆 + 6𝐻_2_𝑂 + 𝑂_2_ → 4γ-𝐹𝑒𝑂𝑂𝐻 + 4𝐻_2_𝑆 (total reaction [3]). Because the discharge capacity of RBM-II was measured by the MFC system, the proton produced by the discharge treatment (reaction [1]) was consumed by the cathode reaction [2], supported that pH was not decreased in RBM-II under the discharged conditions (Fig. 3 A-1). About a charge reaction from lepidocrocite to mackinawite, 4γ-𝐹𝑒𝑂𝑂𝐻 + 4𝐻_2_𝑆 + 8𝑒^-^ → 4𝐹𝑒𝑆 + 2𝑂_2_ + 4𝐻_2_𝑂 + 4𝐻^+^ (reaction [4]) (Fig. S5-B).

Actual reactions were more complicated compared with the predicted reactions because both mackinawite and lepidocrocite were present simultaneously with increase of rechargeable treatments in RBM-II and RCM (Fig. 4). Amorphous iron oxide compound(s), which is a precursor of mackinawite and lepidocrocite (43, 44), was/were detected in RBM-II under charge and discharge conditions (Fig. 5). Furthermore, not all of the sulfide liberated on discharged conditions was used to form mackinawite from lepidocrocite due to diffusion (Fig. 3A-2, B-2), which caused mixture conditions of mackinawite and lepidocrocite. It has been reported that FeS formation can be directly linked to the sulfidation of ferric(hydro)oxides (45). Oxidation of sulfide and subsequent formations of polysulfides and disulfide are key reactants for forming ferrous sulfide minerals via interactions between sulfide and ferric(hydro)oxides (45−47). Previous reports (48−50) supported the importance of sulfur speciation for the form-change of RBM-II. The form-change of CSM was more drastic than that of RBM-II (Fig. 3). It is unknown why the form-change of RBM-II and CSM was different from each other, however, the presence of organic compounds derived from strain HK-II or itself on RBM-II would affect the form-change on rechargeable treatments.

High current densities were observed in all RBM-MFCs at day 0 to day 1 (Fig. 6A) without consumption of lactate as electron donor (Fig. S4), indicating that the current was not produced by strain HK-II but would be produced by charged RBM-II which was mackinawite induced by strain HK-II. RBM-MFCs degraded lactate even under opened circuit conditions and produced high current densities after closed circuit (Fig. 6B and 6C), demonstrated that RBM-II is a useful rechargeable material because of charging electrons at extremely low current density produced from bacterial cells, and indicating the possibility of development of a secondary battery-type MFC. As serious problems of MFCs, it has been pointed out that the current densities were too low to use them for practical applications. Furthermore, MFCs have no mechanism for keeping the degrading activity of organic compounds without current production, because the current production is coupled with organic compounds-degradation, which is the principle of MFCs. A considerably increase of current density and a constant degradation of organic compounds are required for practical use. Electrically conductive nanoparticles have been widely used to improve current density in MFCs (51−53), however, there is no report about a battery type MFC. Our research is first report about a secondary battery-type MFC using biogenic mineral equipped in anode electrodes. Coulombic efficiencies of RBM-MFCs differed from each other (Fig. 6), suggesting that RBM-II equipped in anode electrodes was not sufficiently contacted to the anode surface. It is important for practical application how to assemble anode and RBM-II.

Iron-sulfide(s) are abundant / ubiquitous in subsurface groundwater, sediment and sedimentary rocks (54). These minerals have been studied from biological (55−57) and chemical perspective (48), indicating the relation to the remediation of polluted environments (58−61). These results indicate that microbes and minerals interact closely with each other beyond what was previously general knowledge. One of which is mackinawite due to be produced/induced by biological (57, 58, 62) and chemical reactions (42, 59, 63) and be stable under anaerobic conditions, whereas mackinawite is known as precursor for pyrite (64) and absorbs other metallic ions (65, 66), can be chemically transformed into greigite (49, 67), goethite (α-FeOOH) (48) and leptidocrocite (γ-FeOOH) (50). Amorphous FeS transforms on ageing into makinawite, greigite, pyrrhotite, marcasite, pyrite, and troilite, indicating importance of mackinawite as precursors for other iron sulfides (43, 44), which supports our data shown in this study.

There are electroactive humus and mineral particles in soil and sediment environments, and electroactive bacteria including sulfate-reducers and iron-reducers produce ATP coupled with EET and EEU (67, 68), suggesting that electroactive microbes are relevant to geochemical reactions beyond our consideration. Deep-sea hydrothermal vents, which are environments isolated from solar energy, provide an ideal habitat for chemolithotrophic microbial communities by continuously supplying reductive energy and current via chimneys (23, 29, 69) like a conductive material such as metal (29), resulting in spontaneous and widespread electricity generation around deep-sea hydrothermal fields (23). Conductive consortia, microbial communities, and populations are formed around conductive minerals (70, 71) or via direct interspecies electron transfer (DIET) (72–74). Several bacteria are capable of electrosynthesis by harvesting electrons from conductive materials outside of cells (63, 75). Sulfate reducing bacterium *Desulfovibrio* sp. strain JY forms a conductively methanogenic aggregates with *Methanobacterium* sp. strain YSL via DIET (76). *Shewanella loihica* strain PV-4 self-organizes an electrically conductive network using iron sulfide produced by itself, resulting in the efficient transfer of metabolized electrons (28, 29, 77). These results suggest that biogenic rechargeable minerals such as RBM-II function as electron pool for electron donors and acceptors in microbial ecosystems, which give a cue for understanding the mechanism how slow-growing microbial communities survive in deeply buried sediments where is thought to struggle to obtain cellular maintenance energy (78−80). This expands our understanding of the mechanism how the ubiquity of microbial life even in deep subseafloor environments (78, 81, 82) is maintained irrespective of environments isolated from solar energy.

As conclusion, *Cupidesulfovibrio* sp. strain HK-II induced rechargeable biogenic mineral, RBM-II, which showed incompletely reversible form-change between mackinawite on charge, lepidocrocite on discharge, and amorphous iron oxides on both conditions. The rechargeable reactions of RBM-II were predicted from the results of electrochemical and material science analyses, whereas the rechargeable reactions of CSM remain to be determined. Mackinawite is a ubiquitous mineral in anaerobic environments, suggesting that biogenic and chemically synthesized mackinawites would play roles as electron donors/acceptors for microbial ecosystems not only separated from the solar energy system but also in ordinary anaerobic environments such as paddy fields. Electrical energy acquisition mechanism via rechargeable/conductive minerals such as RBM-II would be one of major survival strategies of microbes because biogenic minerals such as RBM-II would play a role in electron pools as electron donor/acceptor for anaerobes. The EET mechanism of strain HK-II warrant further study as well as analysis of complete genome sequence. Further studies would expand the understanding the overlooked microbial ecosystems and geochemical cycles via electric flow among organic compounds, mineral, and microbes.

## ACKNOWLEDGMENTS

This research was funded in-part by grants KAKENHI (B) 18H03400, KAKENHI (B) 21H03633, and Japan Science and Technology Agency, Crest Grant Number JPMJCR2003.

