## Supplemental figures and tables for "Rechargeable Biomineral Induced by Sulfate Reducing Bacterium *Cupidesulfovibrio* sp. HK-II"

### Supporting Figure S1

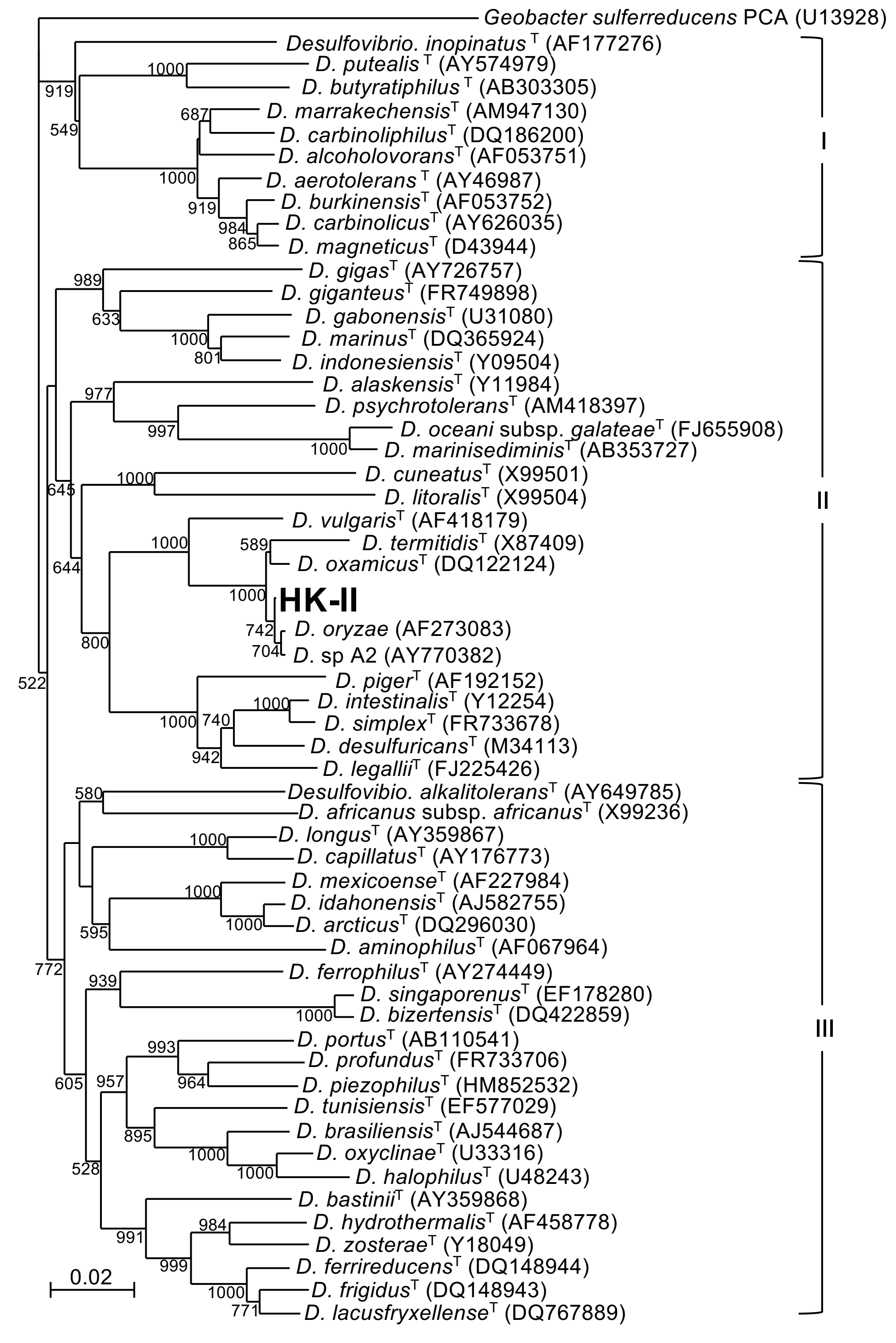


**Phylogenetic analyses of the strain HK-II isolated in this study.**

By homology search of the 16S rRNA gene nucleotide sequence of the strain HK-II, it was shown that the strain HK-II was closely related to *Desulfovibiro oryzae* (99.7% identity, accession number AF273083), identifying the strain HK-II phylogenetically as *Desulfovibiro* sp. strain HK-II. *Desulfovibrio* strains were grouped into three clusters, and the strain HK-II belonged to the cluster II.

### Supporting Figure S2

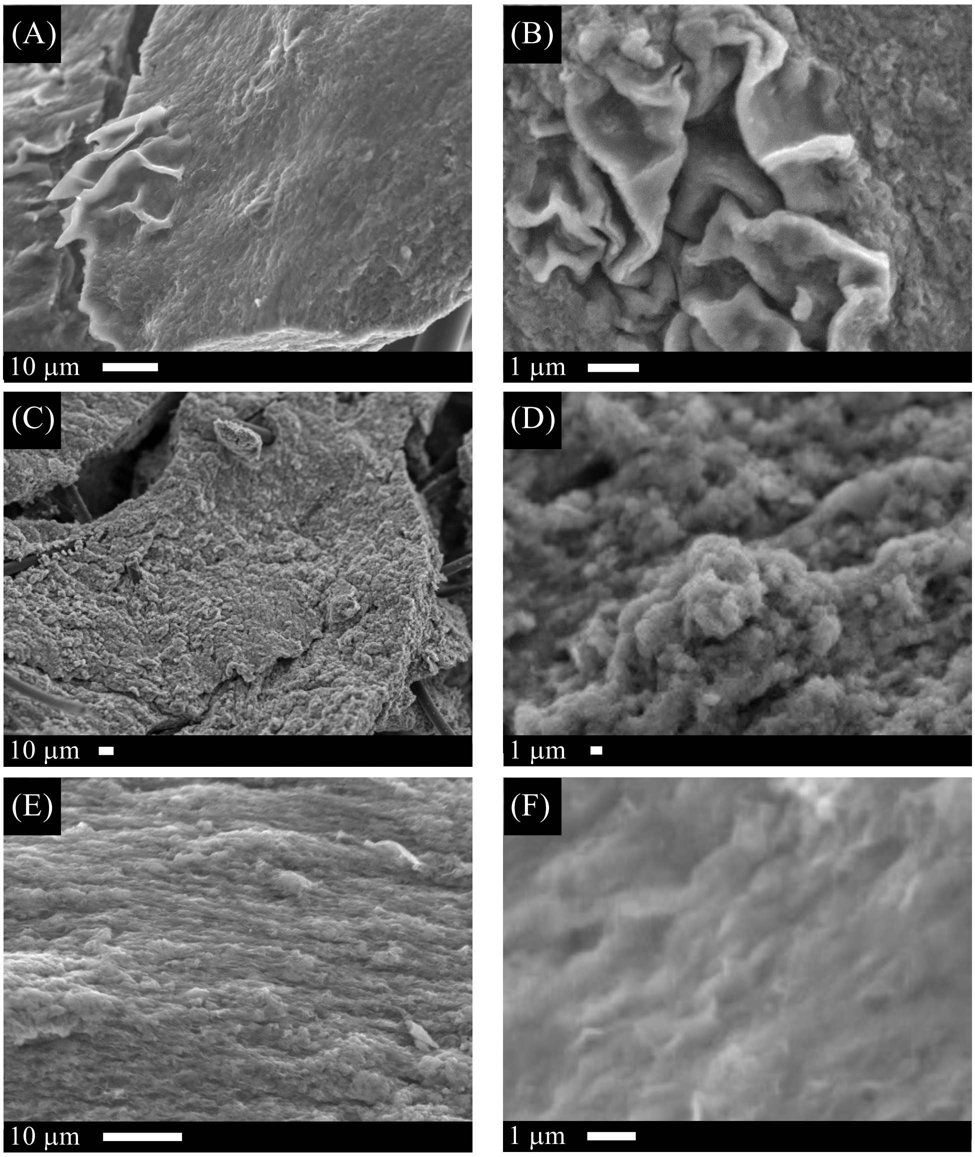


**SEM observation of RBM-II on rechargeable treatments.**

(A and B) RBM-II produced by strain HK-II. Since it is charged, RBM-II is mackinawite. (C and D) RBM-II after the 2nd discharge treatment. Since it is discharged, RBM-II is lepidocrocite. (E and F) RBM-II after the 2nd charge treatment. Since it is charged, RBM-II is mackinawite. White bars are size markers (10 µm or 1 µm).

### Supporting Figure S3

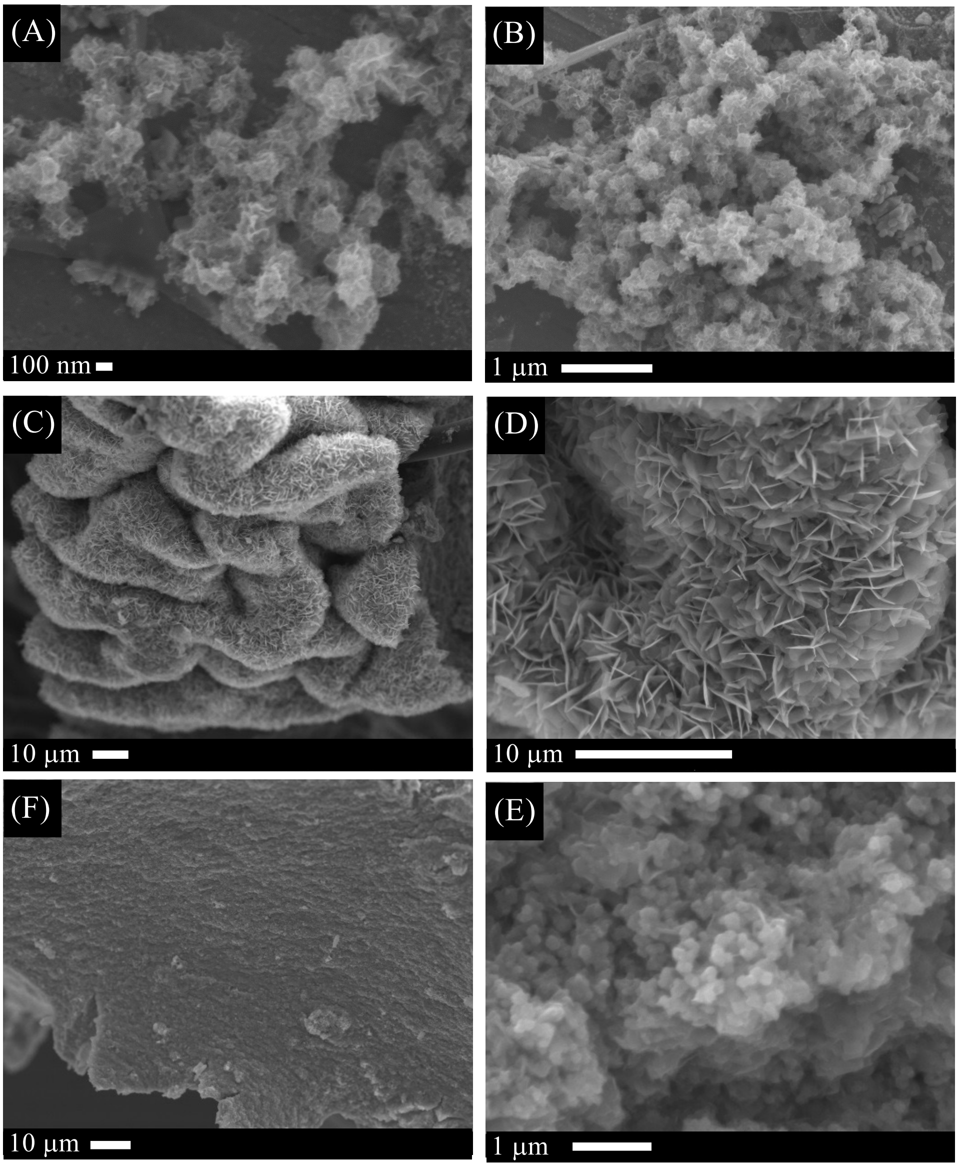


**SEM observation of CSM on rechargeable treatments.**
(A and B) CSM produced by chemical reactions. Since it is charged, CSM is mackinawite. (C and D) CSM after the 2nd discharge treatment. Since it is discharged, CSM is lepidocrocite. (E and F) CSM after the 2nd charge treatment. Since it is charged, CSM is mackinawite. White bars are size markers (10 μm or 1 μm).

### Supporting Figure S4

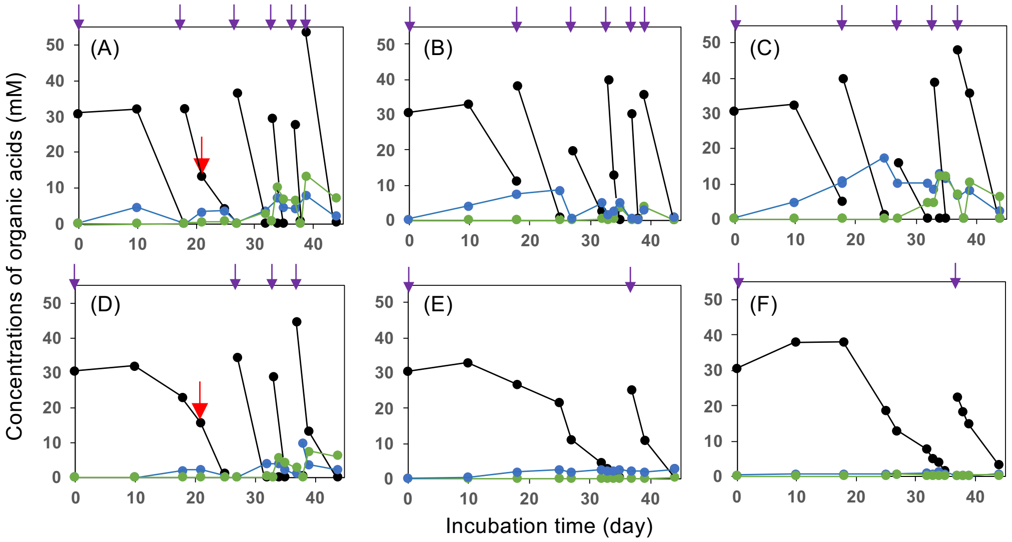


**Concetrations of organic acids in MFCs.** (A-C): RBM-MFCs. (A); RBM-MFC1, (B); RBM-MFC2, (C); RBM-MFC3, (D-F): Control-MFCs. (D); control-MFC1, (E); control-MFC2, (F); control-MFC3. Black circles; lactate, blue circulars; acetate, green circulars; propionate. Red arrows donate the time of addition of sulfate. Purple arrows donate the time of addition of sodium lactate.

### Supporting Figure S5

#### Rechargeable reactions of RBM-II

##### A) Discharge reaction

$4FeS + 8H_2O \rightarrow 4\gamma\text{-}FeOOH + 4H_2S + 4H^+ + 4e^-$ (reaction on the anode [1])
$4H^+ + O_2 + 4e^- \rightarrow 2H_2O$ (reaction on the cathode [2])

total reaction was as follows;

$4FeS + 6H_2O + O_2 \rightarrow 4\gamma\text{-}FeOOH + 4H_2S$ (total reaction [3]).

Discharge reaction on the anode was deduced as follows. The reaction consists of 3 steps at least.

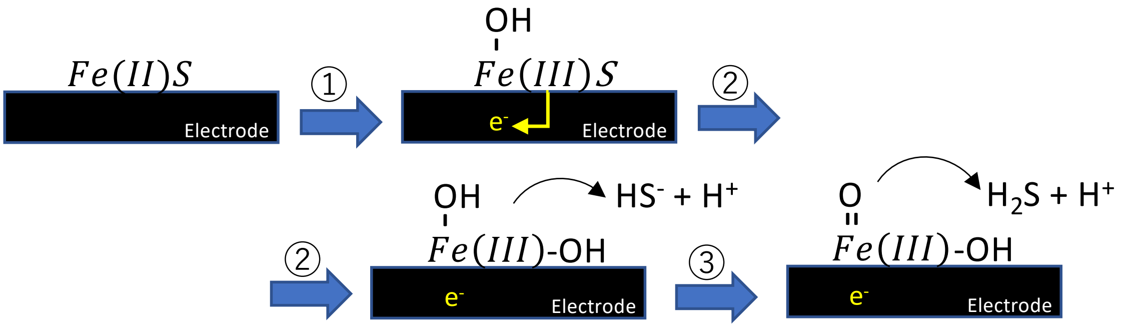


##### B) Charge reaction $4\gamma\text{-}FeOOH + 4H_2S + 8e^- \rightarrow 4FeS + 2O_2 + 4H_2O + 4H^+$

Charge reaction was deduced as follows. The reaction consists of 4 steps at least.

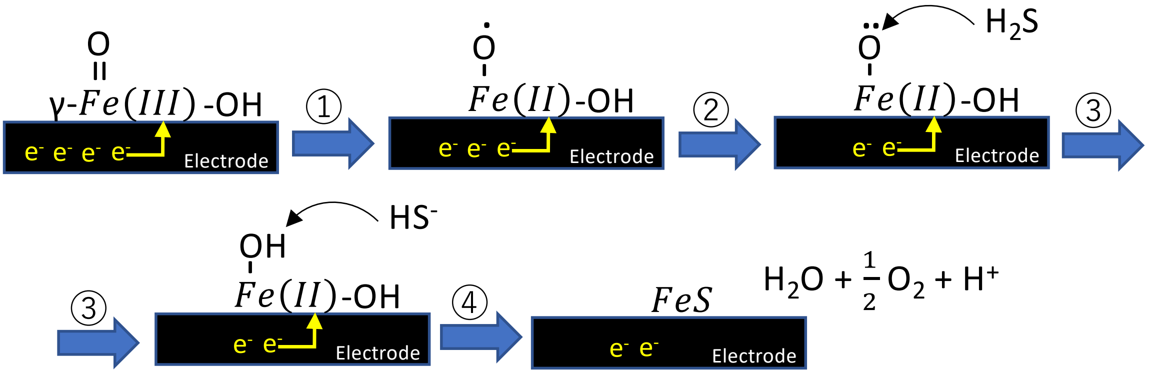

Table S1. Crystallite size of RBM-II, and CSM

| Plate | Two-theta [°] | crystallite size [nm] | |
|---|---|---|---|
| | | RBM-II | CSM |
| 001 | 17.6 | 6.2 | 8.0 |
| 101 | 30.1 | 6.3 | 11.5 |
| 111 | 39.0 | 12.0 | 12.5 |
| 200 | 49.5 | 12.3 | 9.6 |
| Average | | 9.2±1.5 | 10±0.9 |

Table S2. Bindig energies for Fe ($2p^{3/2}$), S(2p), and O(1s) peaks in comparison with fitted peaks in Fig 4.

| | Binding energy (ev) | Species | References |
|---|---|---|---|
| Fe ($2p^{3/2}$) | 707.1 | Fe(II)-S | Pratt et al., 1994 (39) |
| | 707.3 | Fe(II)-S | Herbert et al. 1998 (36) |
| | 707.5 | Fe(II)-S | Jones et al., 1992 (37) |
| | 709.0 | Fe(II)-O | Thomas et al., 1998 (38) |
| | 709.5 | Fe(II)-O | McIntyre and Zetaruk 1977 (35) |
| | 709.7 | Fe(III)-S | Thomas et al., 1998 (38) |
| | 711.0 | Fe(III)-O | Thomas et al., 1998 (38) |
| | 711.6 | Fe(III)-O | Mullet et al., 2002 (42) |
| | 712.0 | Fe(III)-O | Thomas et al., 1998 (38) |
| | 714.0 | Fe(III)-O | Thomas et al., 1998; Pratt et al., 1994 (38, 39) |
| S(2p) | 160.95 | $S^{2-}$ | Herbert et al., 1998 (36) |
| | 161.25 | $S^{2-}$ | Pratt et al., 1994 (39) |
| | 161.3 | $S^{2-}$ | Pratt et al., 1994, Mullet et al., 2002 (39, 42) |
| | 161.4 | $S^{2-}$ | Thomas et al., 1998 (38) |
| | 162.2 | $S_2^{2-}$ | Herbert et al. 1998 (36) |
| | 162.25 | $S_2^{2-}$ | Herbert et al. 1998 (36) |
| | 162.5 | $S_2^{2-}$ | Mycroft et al., 1990 (40) |
| | 163.15 | $S_n^{2-}$ | Herbert et al. 1998 (36) |
| | 163.4 | $S_n^{2-}$ | Thomas et al., 1998 (38) |
| | 164.0 | $S_8$ | Thomas et al., 1998 (38) |
| O(1s) | 529.5 | $O^{2-}$ | Mullet et al., 2002 (42) |
| | 529.8 | $O^{2-}$ | Jones et al., 1992 (37) |
| | 530.0 | $O^{2-}$ | Ferris et al., 1989 (41) |
| | 530.2 | O (Lepidocrocite) | McIntyre and Zetaruk 1977 (35) |
| | 531.3 | $OH^-$ | Mullet et al., 2002 (42) |
| | 531.4 | $OH^-$ (Lepidocrocite) | McIntyre and Zetaruk 1977 (35) |
| | 531.5 | $OH^-$ | Herbert Jr. et al. 1998 (36) |
| | 532.52 | adsorbed $H_2O$ | Herbert Jr. et al. 1998 (36) |
| | 532.6 | adsorbed $H_2O$ | Pratt et al., 1994 (39) |
| | 533.58 | other $H_2O$ | Herbert Jr. et al. 1998 (36) |
